# Vacuolar iron stores gated by NRAMP3 and NRAMP4 are the primary source of iron in germinating seeds

**DOI:** 10.1101/306894

**Authors:** Emma L Bastow, Vanesa S Garcia de la Torre, Andrew E Maclean, Robert T Green, Sylvain Merlot, Sebastien Thomine, Janneke Balk

## Abstract

During seed germination, iron (Fe) stored in vacuoles is exported by the redundant NRAMP3 and NRAMP4 transporter proteins. A double *nramp3 nramp4* mutant is unable to mobilize Fe stores and does not develop in the absence of external Fe. We used RNA sequencing to compare gene expression in *nramp3 nramp4* and wild type during germination and early seedling development. Even though sufficient Fe was supplied, the Fe-responsive transcription factors *bHLH38, 39, 100* and *101* and their downstream targets *FRO2* and *IRT1* mediating Fe uptake were strongly upregulated in the *nramp3 nramp4* mutant. Activation of the Fe deficiency response was confirmed by increased ferric chelate reductase activity in the mutant. At early stages, genes important for chloroplast redox control (*FSD1, SAPX*), Fe homeostasis *(FER1, SUFB)* and chlorophyll metabolism (*HEMA1, NYC1*) were downregulated, indicating limited Fe availability in plastids. In contrast, expression of *FRO3*, encoding a ferric reductase involved in Fe import into the mitochondria, was maintained and Fe-dependent enzymes in the mitochondria were unaffected in *nramp3 nramp4*. Together these data show that a failure to mobilize Fe stores during germination triggered Fe deficiency responses and strongly affected plastids but not mitochondria.

## INTRODUCTION

Fe is essential for multiple pathways in plants, therefore the uptake, storage, redistribution and recycling of Fe are highly regulated (Connorton et al., 2017; Jeong et al., 2017). Transcriptional regulation during Fe deficiency has been extensively studied in seedlings grown on agar plates or in adult plants grown in hydroponic conditions, leading to the identification of many genes involved in Fe homeostasis (Buckhout et al., 2009; Colangelo et al., 2004; Dinneny et al., 2008; Mai et al., 2016; Rodríguez-celma et al., 2013). However, the gene networks are very complex, as transcriptional changes occurring in different cell types are usually stacked together, and mechanisms to restrict nutrient use, release Fe from storage and increase uptake are induced simultaneously.

During germination and early development, the seedling relies primarily on its Fe stores before it has developed a root to take up Fe from the environment. While some seeds store Fe in the form of ferritin (Briat et al., 2010), oilseeds such as *Arabidopsis thaliana* store Fe in vacuoles of the root endodermis and around the provasculature in the cotyledons (Kim et al., 2006; Roschzttardtz et al., 2009). The *Vacuolar Iron Transporter VIT1* is expressed during seed development to enable Fe storage into endodermal vacuoles (Kim et al., 2006). Fe is exported from the vacuoles by NRAMP3 and NRAMP4, two redundant divalent cation transporters belonging to the family of Natural Resistance-Associated Macrophage Proteins (Lanquar et al., 2005). *NRAMP3* and *NRAMP4* are highly expressed in the first few days after sowing. Total Fe content and localization are unaffected in mature seeds of *nramp3 nramp4* double mutants (Ramos et al., 2013). However, when grown in medium lacking Fe, *nramp3 nramp4* mutants have short roots, chlorotic leaves and their growth is arrested (Lanquar et al., 2005). Development and greening of *nramp3 nramp4* seedlings may be restored by providing external Fe in the medium.

Organelles such as mitochondria and chloroplasts have a high demand for Fe as they contain electron transport chains and metabolic pathways that require numerous Fe cofactors. Synthesis of iron-sulfur (FeS) clusters and haem is therefore essential for these organelles (Lill et al., 2012; Balk & Schaedler 2014). In photosynthetically active leaf cells, over 80% of the cellular Fe is localized in chloroplasts (Lanquar et al., 2010; Shingles et al., 2002). Photosystems I and II, cytochrome *b_6_f*, ferredoxins and Fe superoxide dismutase (FeSOD) are the main proteins that utilise Fe cofactors. During germination, most plant species are heterotrophic, relying entirely on energy stores to make ATP. The bulk of ATP is produced by the mitochondria, which become bioenergetically active immediately upon hydration (Paszkiewicz et al., 2017). Mitochondrial respiration is highly dependent on Fe enzymes, such as respiratory chain complexes I - IV, aconitase and ferredoxins. FeS clusters are also synthesized in the cytosol for enzymes such as cytosolic aconitase, aldehyde oxidases and DNA repair enzymes in the nucleus (Balk & Schaedler 2014).

Using a transcriptomic approach, we have compared global gene expression patterns between an *nramp3 nramp4* double mutant and wild-type Arabidopsis during early development. We analysed the transcriptional differences of 1-day old (imbibition), 3-day old (radicle emergence) and 8-day old (green cotyledon) plants alongside protein levels and enzymatic activities to gain insight into regulation of Fe-dependent processes at the transcriptional and post-transcriptional level. We show that during early development, the *nramp3 nramp4* mutant triggers a typical Fe deficiency response even in the presence of Fe in the medium. Transcription of many genes for chloroplast functions were decreased in *nramp3 nramp4*. In contrast, only a small number of genes encoding mitochondrial proteins were differentially expressed and essential functions of mitochondria were maintained.

## RESULTS AND DISCUSSION

### A limited set of genes is differentially expressed in germinating *nramp3 nramp4* seedlings supplied with sufficient Fe

To investigate differential gene expression in early seedling development between *nramp3 nramp4* and wild type, seeds were harvested from plants grown side-by-side in a controlled environment and germinated in liquid medium containing 50 μM Fe. Under these conditions, development of mutant and wild-type seedlings was similar except that cotyledons were slightly chlorotic in *nramp3 nramp4* at 8 days (Figure 1). Plant material was collected after 24 h imbibition (growth stage 0.10 according to Boyes et al., 2001), after 72 h / 3 days upon radical emergence (growth stage 0.50), and after 8 days when cotyledons were fully expanded (growth stage 1.00). RNA was extracted for preparation of mRNA libraries which were sequenced using Illumina technology. Between 35 and 42 million reads were obtained for three independent biological replicates of wild-type and *nramp3 nramp4* at each growth stage and mapped to the *Arabidopsis* TAIR10 genome (Table S1; Figure S2). Combining all time points and using a Fold-Change cut off > 3.0 (*p* < 0.05), only 302 genes were differentially expressed between wild-type and *nramp3 nramp4* plants out of a total of 18,493 expressed genes (Figure 2; Table S2). As expected, the number of RNA reads corresponding to *NRAMP3* was decreased in *nramp3 nramp4* compared to wild type at day 1 and 3 (Figure 3). The distribution of reads along the *NRAMP3* gene indicates that transcription is initiated downstream of the T-DNA insertion, resulting in a transcript lacking the first ~ 100 nucleotides of coding sequence and most likely a non-functional protein (Figure S1). At day 8, wild-type expression of *NRAMP3* is very low, and therefore not different from the double mutant. For *NRAMP4*, very few RNA reads map to the coding sequence downstream of the T-DNA insertion and little full-length transcript is produced (Figure S1). However, RNA reads upstream of the T-DNA insertion may give the false impression that *NRAMP4* is expressed at almost similar levels in mutant and wild type at day 3 (Figure 3).

**Figure 1.**
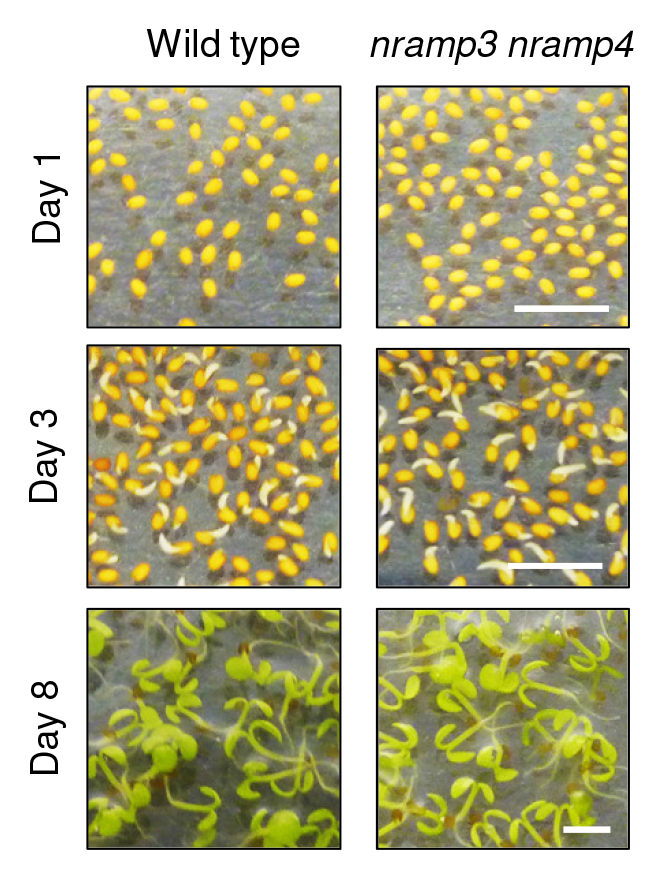
Germination of the *nramp3 nramp4* double mutant in the presence of Fe. Wild type and *nramp3 nramp4* after imbibition (day 1), radical emergence (day 3) and cotyledon emergence (day 8). Scale bar 2 mm.

**Figure 2.**
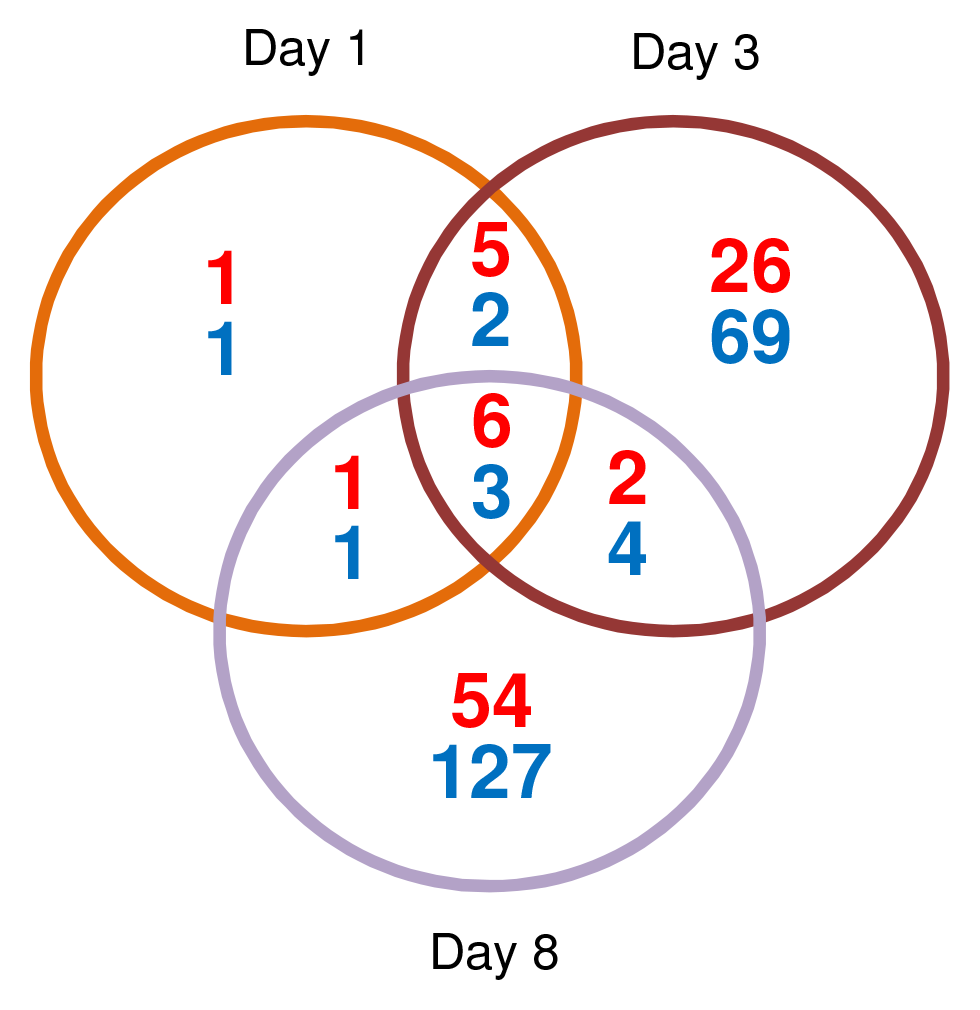
Number of differentially expressed genes in *nramp3 nramp4* compared to wild type during early development. Upregulated (red) and downregulated (blue) genes in *nramp3 nramp4* compared to wild type at 3 stages of germination. The total number of expressed genes analysed was 18,493, of which only 302 genes were differentially expressed using FC > 3, n = 3, *p* < 0.05 for each time point.

**Figure 3.**
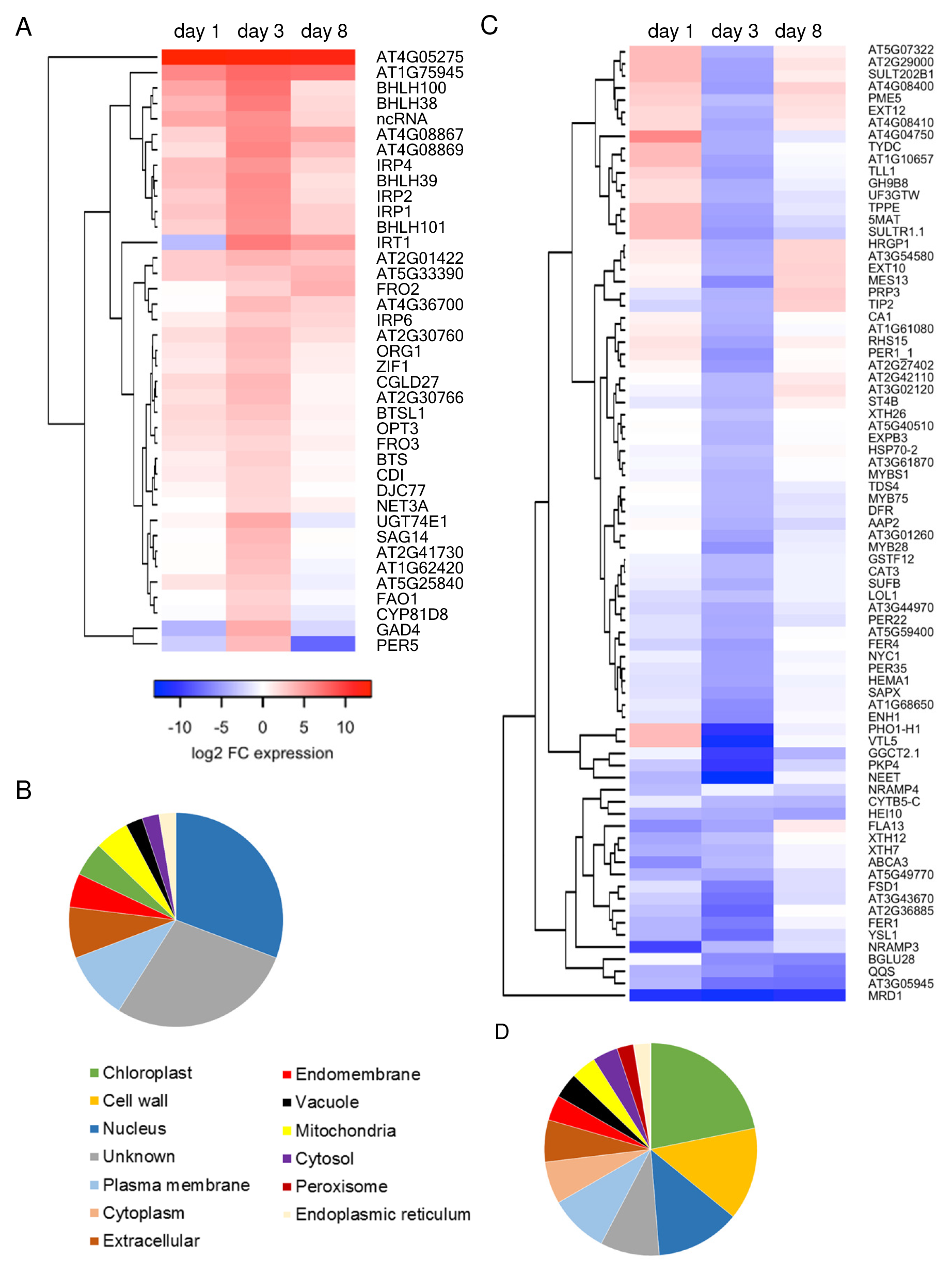
Differentially expressed genes in *nramp3 nramp4* (FC > 3) and predicted protein localization. A, Heatmap of transcript levels with >3-fold upregulation in *nramp3 nramp4* compared to wild type (*p* < 0.05). B, Predicted subcellular localisations of the 39 proteins encoded by the upregulated genes. C, Heatmap of transcript levels with >3-fold downregulation in *nramp3 nramp4* compared to wild type (*p* < 0.05). D, Predicted subcellular localisations of the 78 proteins encoded by downregulated genes.

Comparing *nramp3 nramp4* and wild-type plants at day 1, a total of 20 transcripts were differentially expressed (Figure 2). By day 3, the total number of differentially expressed transcripts was 117, of which 16 were common between day 1 and day 3 plants. At day 8, the number of differentially expressed transcripts had increased to 198 but this set had little overlap with the 3-day time point (183 non-common transcripts). For several genes downregulated at day 3, expression was recovered at day 8. This suggests that secondary responses are induced in 8-day old *nramp3 nramp4* plants, since *NRAMP3* and *NRAMP4* expression levels have declined in wild type at that stage (see above, Lanquar et al., 2005). We therefore focussed on the 3-day time point, corresponding to the highest expression level of *NRAMP3* and *NRAMP4*, for further comparative analysis of upregulated (Figure 3A and Table S3) and downregulated genes (Figure 3C and Table S4). The differentially expressed genes were classified according to cellular localization of the gene products, which revealed that predicted nuclear proteins are relatively overrepresented in the upregulated genes, whereas in the downregulated genes chloroplast and cell wall proteins are overrepresented (Figure 3C, D).

### The Fe deficiency response is induced in *nramp3 nramp4* seedlings germinating in the presence of exogenous Fe

The upregulated genes include four basic helix-loop-helix (bHLH) transcription factors that control activation of the Fe deficiency response: *bHLH38, bHLH39, bHLH100* and *bHLH101*. Increased transcript levels of *bHLH38* in *nramp3 nramp4* relative to wild type was confirmed by qRT-PCR at all three time-points (Figure 4A). bHLH38 and bHLH39 have been shown to form a dimer with FIT (bHLH29) and directly activate transcription of the *Iron-Regulated Transporter IRT1* and the *Ferric Reductase Oxidase FRO2* (Yuan et al., 2008; Wang et al., 2013). Although *FIT* expression was not altered in the mutant, *IRT1* and *FRO2* were upregulated in *nramp3 nramp4* at the 3-day and 8-day time points. *FRO2* transcript levels were increased ~ 16-fold in 8-day-old *nramp3 nramp4* seedlings compared to wild type (Supplemental Table S3), in agreement with RT-qPCR analysis (Figure 4B). Accordingly, ferric reductase activity displayed a 2-fold increase at the same time point (Figure 4C).

**Figure 4.**
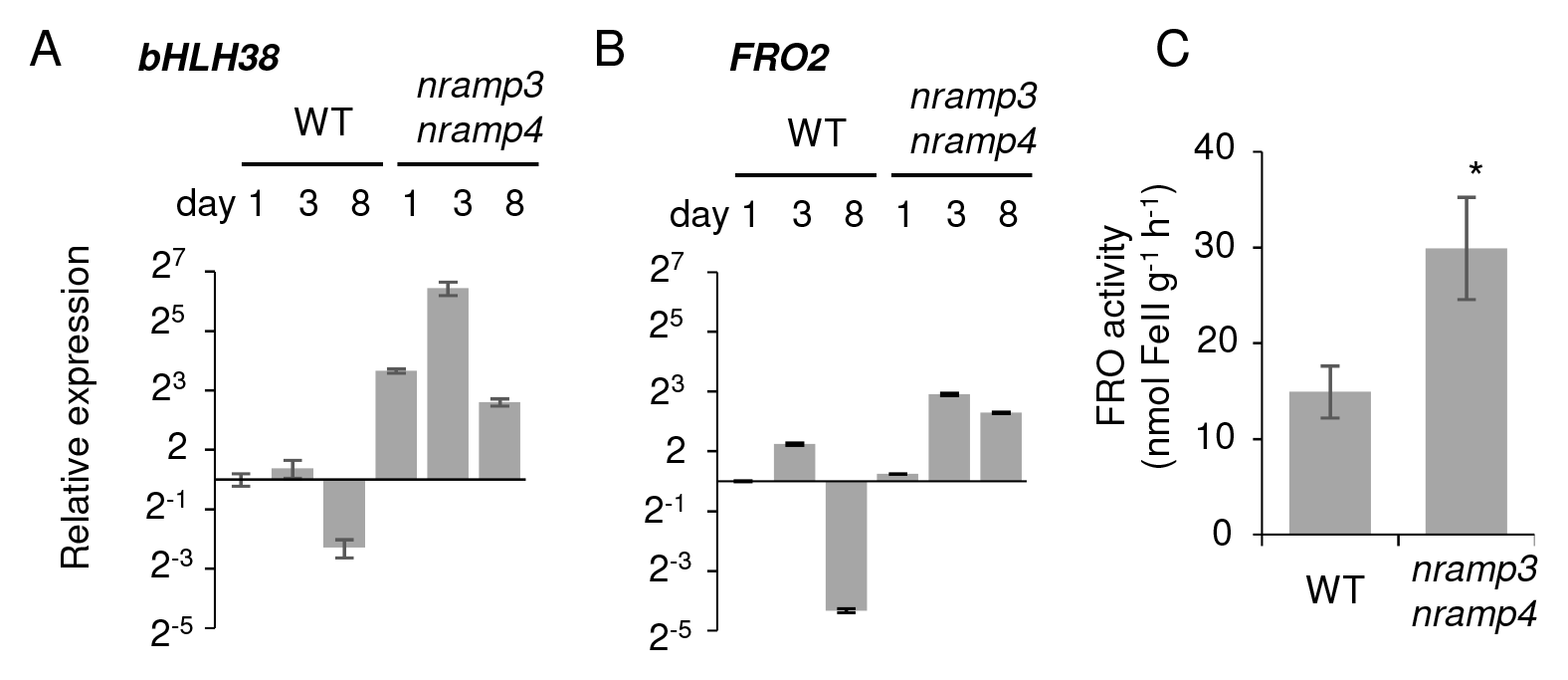
*nramp3 nramp4* seedlings activate the Fe deficiency response. A, RT-qPCR of *bHLH38*. B, RT-qPCR of *FRO2*. For A and B, values are the mean of 3 biological replicates ± SE. C, Ferric reductase activity of 8-day-old wild-type and *nramp3 nramp4* seedlings, measured by the formation of Fe(II)-ferrozine in a spectrophotometric assay at 562 nm. Values are the mean of 3 biological samples of pooled seedlings ± SD, ^*^*p* < 0.05.

Other genes belonging to the core set of the ferrome (Buckhout et al., 2009; Mai et al., 2016) are also upregulated in *nramp3 nramp4* during germination. These genes encode the following proteins: the oligopeptide transporter OPT3 required for Fe loading into the phloem; the nuclear protein kinase ORG1 and the uncharacterized Iron-Regulated Proteins IRP1, IRP2, IRP4 and IRP6 (Rodríguez-celma et al., 2013). The E3 ubiquitin-protein ligases BRUTUS (BTS) and BTSL1, negative regulators of Fe homeostasis (Hindt et al., 2017; Kobayashi et al., 2013; Selote et al., 2015) are also upregulated in *nramp3 nramp4*. The vacuole-located ZIF1 was upregulated and its role in increasing the concentration of the metal chelator nicotianamine (NA) in the vacuole (Haydon et al., 2012) suggests an attempt to mobilize vacuolar Fe as an Fe-NA complex in the *nramp3 nramp4* mutant. It is noteworthy that many genes previously shown to participate in the Fe deficiency response are not upregulated in germinating *nramp3 nramp4* even though their expression is detected. This is the case for *F6’H1* and *PDR9* that allow the release of coumarins in the rhizosphere to mobilize Fe (Tsai & Schmidt, 2017), *NRAMP1* for low affinity Fe uptake as well as *MTP3, IREG2* and *MTP8*that sequester excess heavy metal imported by IRT1 (Thomine & Vert, 2013; Castaings et al., 2016). This suggests that the transcriptional Fe deficiency response is modulated according to the developmental stage.

Taken together, the RNA-seq data, qRT-PCR and the ferric chelate reductase activity measurements show that Fe deficiency responses are activated in *nramp3 nramp4* even in Fe-sufficient conditions. This indicates that at early stages of development, Arabidopsis seedlings rely on their Fe stores rather than the environment to acquire sufficient Fe. The induction of the Fe deficiency response including *IRT1* allows the mutant to overcome the defect in vacuolar export.

### Iron supply to plastids is delayed when vacuolar Fe cannot be retrieved

Many downregulated genes in *nramp3 nramp4* (17 out of 78) encode proteins predicted to localize to the chloroplast (Figure 3D). Expression of two ferritin genes, *FER1* and *FER4* was decreased in 1-day-old and 3-day-old plants, but similar to wild type at 8 days (Figure 3C). This pattern of expression was confirmed by qRT-PCR of *FER1* (Figure 5A), and immunodetection of ferritin protein (Figure 5B). Thus, in the absence of Fe mobilization from the vacuoles ferritin expression is strongly decreased, suggesting that the *nramp3 nramp4* seedlings limit Fe availability to the developing plastids as an “Fe sparing” strategy.

**Figure 5.**
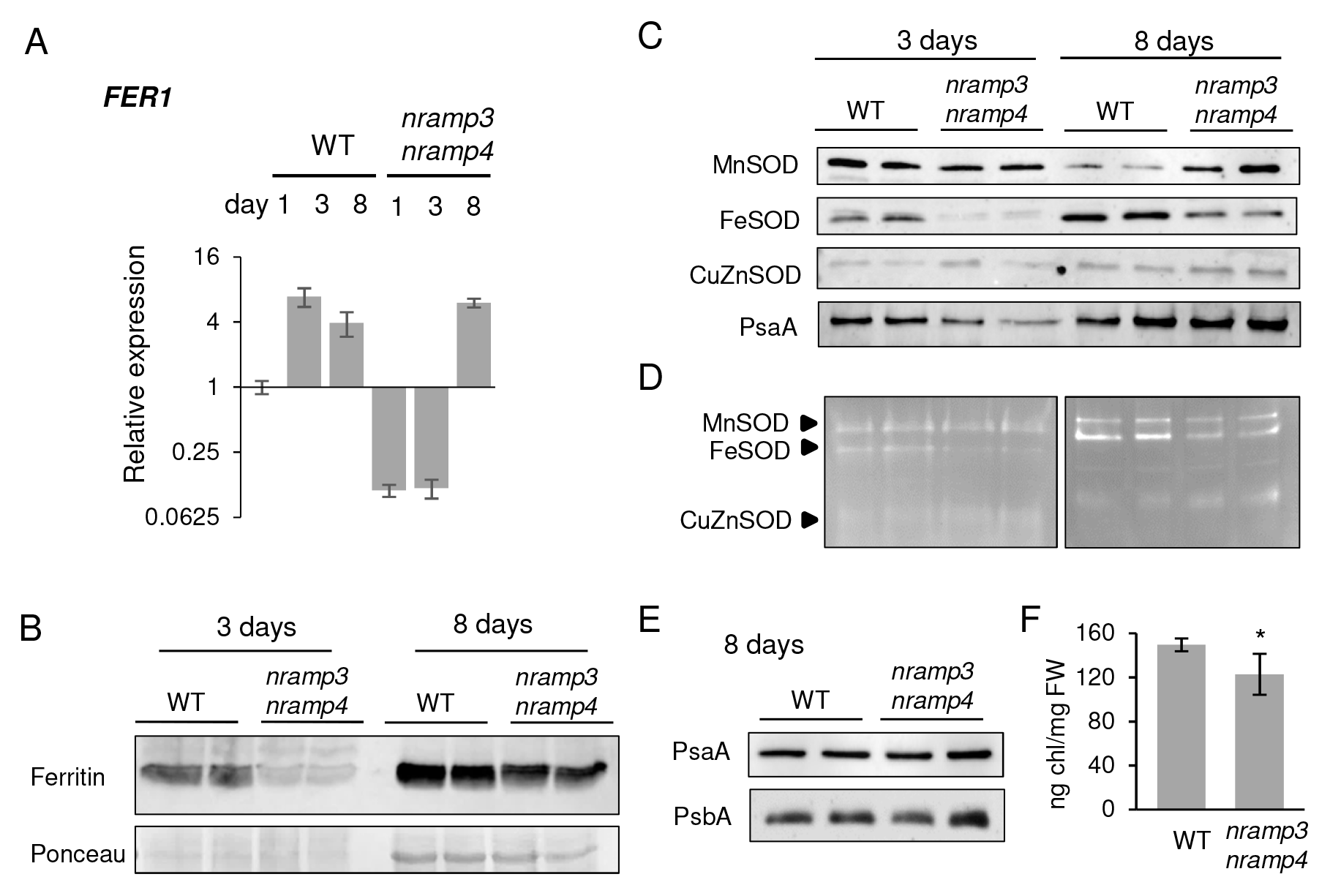
Plastid-localized Fe proteins are decreased in *nramp3 nramp4* A, qRT-PCR of *FER1*. Values are the mean of 3 biological replicates ± SE. B, Ferritin protein levels of 3-day and 8-day-old plants by Western blot analysis. C, Western blot analysis of Superoxide Dismutase (SOD) proteins in extracts from 3-day and 8-day old wild-type (WT), and *nramp3 nramp4* seedlings. Immunodection of PsaA served as a control for equal loading. D, SOD activities revealed by nitro blue tetrazolium, which appears as negative staining, of plant extracts as in (C). E, Western blot analysis of Photosystem I and II subunits in 8-day-old plants. F, Total chlorophyll content in 8-day-old plants measured in a spectrophotometric assay at 645 nm and 662 nm. Values are the mean ± SD (n = 4), ^*^*p* < 0.05, Student’s t-test.

Two genes involved in tetrapyrrole metabolism, *HEMA1* and *NYC1*, are also strongly downregulated in *nramp3 nramp4* (Figure 3C). *HEMA1* encodes glutamyl-tRNA reductase which catalyses the NADPH-dependent reduction of glutamyl-tRNA to glutamate 1- semialdehyde in the first step in tetrapyrrole biosynthesis required for the production of both haem and chlorophylls (Kobayashi et al., 2016). Accordingly, we measured a slight decrease in total chlorophyll content in the mutant in Fe-sufficient conditions (Figure 5F). *NYC1* encodes chlorophyll *b* reductase required for degradation of chlorophyll *b* (Tanaka et al., 2011). Coordinated downregulation of *HEMA1* and *NYC1* was previously observed in Fe deficient leaves (Rodríguez-celma et al., 2013). In contrast, *CGLD27*, a highly conserved gene associated with carotenoid-xanthophyll metabolism involved in protection against excess light stress, was upregulated in *nramp3 nramp4* (Urzica et al., 2012; Rodriguez-Celma et al 2013). Plastids contain the so called SUF pathway for FeS cluster assembly, consisting of 6 proteins which are evolutionary conserved with cyanobacteria and most alpha-proteobacteria. In *nramp3 nramp4* seedlings, the expression of *SUFB* is decreased at 1 and 3 days (Figure 3C). It has been noticed before that *SUFB* is repressed under Fe deficiency whereas other *SUF* genes do not respond to Fe (Balk & Schaedler, 2014). SUFB is a subunit of the FeS cluster scaffold and essential for all plastid-localized FeS proteins (Hu et al., 2017). Depletion of SUFB leads to strongly decreased levels of Photosystem I (PSI), which binds 3 [4Fe-4S] clusters on the PsaA, PsaB and PsaC subunits. However, the level of subunit PsaA of PSI was remarkably stable at 3 and 8 days in the mutant, in agreement with RNA-seq data showing strong expression at all stages of germination. This suggests that PsaA protein is stable without FeS cofactor. PsbA of PSII could not be detected in wild type or *nramp3 nramp4* at 3 days (Figure 5C and not shown). At 8 days, PsaA and PsbB levels were similar in *nramp3 nramp4* and wild type (Figure 5E), when *SUFB* expression was back to wild-type levels (Figure 3C). Presumably, at this stage the mutant seedlings had acquired enough Fe to synthesize FeS clusters and provide PSI with its FeS cofactors. Of the many FeS proteins in plastids, only the stroma-localized [2Fe-2S] protein NEET (Su et al., 2013) was transcriptionally downregulated at day 1 and 3, but not at day 8.

Interestingly, transcripts of genes encoding Fe-binding proteins involved in oxidative stress responses were also decreased. For example, downregulation of *ENH1, SAPX*and *FSD1* that encode rubredoxin, stromal ascorbate peroxidase and FeSOD, respectively, was observed. At the post-translational level, we observed a decrease in FeSOD protein level (Figure 5C) correlating with decreased FeSOD activity (Figure 5D) in both 3- and 8-day-old *nramp3 nramp4* plants. Interestingly, the protein level of MnSOD, which is located in the mitochondria, was increased in 8-day-old mutant seedlings relative to wild type, but there was no difference in MnSOD activity between the 2 genotypes. The protein levels and activity of CuZnSOD were similar in wild-type and *nramp3 nramp4*. Knock-out mutants of *FSD1* have no phenotype, indicating that in plastids CuZnSOD can fully compensate for the lack of FeSOD (Pilon et al., 2011).

### Iron-dependent respiratory complexes in the mitochondria are not affected in germinating *nramp3 nramp4* seeds

Only five genes encoding proteins with either confirmed or predicted mitochondrial localization are differentially expressed in *nramp3 nramp4* at the 3-day time point (Figure 3A, C). The mitochondrial ferric reductase 3 (*FRO3*) was upregulated (Figure 3A), suggesting that mitochondria continue to import Fe (Jain et al., 2013). *MIT1* and *MIT2*, homologs of the well-characterized Mitochondrial Iron Transporter in other species (Bashir et al., 2011) were not differentially expressed, but they generally do not respond to Fe deficiency (Balk & Schaedler, 2014).

To investigate if Fe-binding proteins in the mitochondria were affected post-transcriptionally, we analysed the levels of respiratory complex I, II and III. Complex I binds 8 FeS clusters (22 Fe in total), complex II binds 3 FeS clusters (10 Fe) and complex III binds 4 haem cofactors and one Fe_2_S_2_ cluster (6 Fe). Mitochondria were purified from 3-day-old seedlings and subjected to Blue Native-Poly Acrylamide Gel Electrophoresis to resolve the large membrane complexes. Total protein was stained with Coomassie Brilliant Blue, which showed similar levels of complex I, complex V and complex III in *nramp3 nramp4* and wild type (Figure 6A). Complex II is not clearly visible using Coomassie staining, but its activity can be detected ingel using succinate as substrate and a chromogenic electron acceptor. This showed that complex II activity was not affected in the *nramp3 nramp4* mutant (Figure 6B, lower panel). A similar in-gel staining method specific for Complex I, using NADH as a substrate and electrons passing through only part of the complex, confirmed there was no decrease in complex I levels in *nramp3 nramp4* (Figure 6B, top panel). Our findings contrast with the decrease in complex I that has been observed in roots of cucumber seedlings grown hydroponically without Fe (Vigani et al., 2009) suggesting that priority for Fe allocation may differ according to the organ or the developmental stage. To investigate proteins involved in FeS cluster assembly, we probed total cell extracts from 3-day-old wild-type and *nramp3 nramp4* seedlings for NFU4 and NFU5, using protein blot analysis. The levels of the two NFU proteins were similar in mutant and wild type (Figure 6C). Taken together these data suggest that mitochondria are protected from Fe deficiency during the early stages of growth, either because they have autonomous Fe stores or because Fe is prioritized to this organelle due to its essential function during germination.

**Figure 6.**
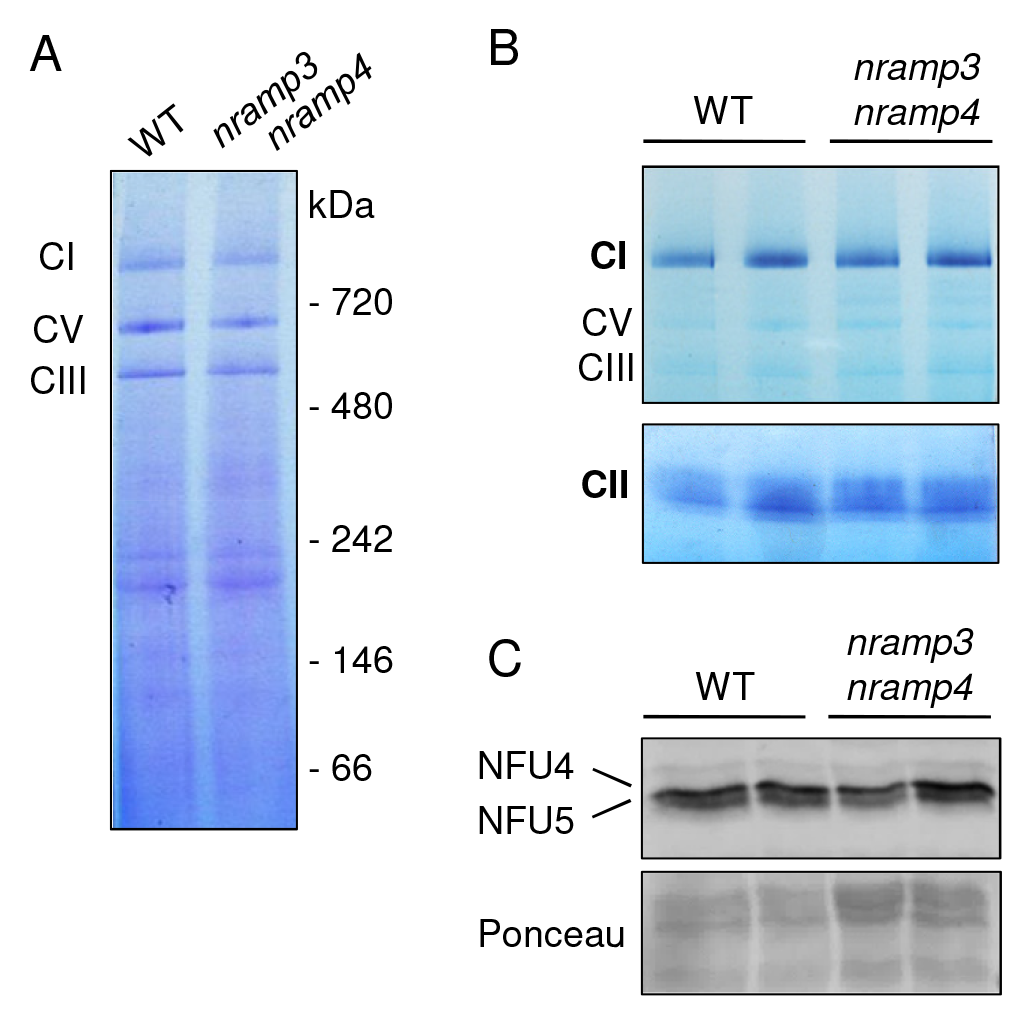
Mitochondrial Fe-dependent enzymes are maintained in *nramp3 nramp4*. A, BN-PAGE analysis of mitochondrial proteins (10μ g) that were isolated from 3-day-old seedlings. Mitochondrial complexes I, V (CV) and III (CIII) were stained with Coomassie Brilliant Blue. B, Activity staining of mitochondrial proteins (25 |jg) from 3- day-old seedlings, separated by BN-PAGE. Complex I (CI) and complex II (CII) activities were visualised using NADH and succinate respectively as electron donor and the colorimetric electron acceptor nitro blue tetrazolium. C, Western blot analysis with antibodies against NFU4 and NFU5 proteins. Ponceau stain was used as a loading control.

### Fe limitation impacts Fe-dependent enzymes in other cellular compartments

Outside of plastids and mitochondria, enzymes that require Fe for function were also affected in *nramp3 nramp4* seedlings. For instance, transcription of *CAT3* was downregulated in *nramp3 nramp4* (Figure 3C, D). CAT3 is one of three catalase isoforms in the peroxisome involved in oxidative stress responses. Accordingly, catalase protein levels were decreased in 3-day-old *nramp3 nramp4*, correlating with decreased catalase activity (Figure 7A, B). Catalase depends on a haem cofactor for activity, therefore the downregulation of tetrapyrrole biosynthesis (see above) is likely to have an impact on haem enzymes throughout the cell. The enzyme aconitase depends on a Fe_4_S_4_ cofactor. During germination, aconitase is highly upregulated to mobilize storage lipids *via* the glyoxylate cycle. This is due to specific induction of the *ACO3* gene, of which the gene product is localized in the cytosol at this developmental stage (Hooks et al., 2014). Although transcription of *ACO1, ACO2 and ACO3* and aconitase protein levels were unaffected in *nramp3 nramp4*, aconitase activity was strongly decreased (Figure 7C, D). Iron limitation therefore impacts cytosolic aconitase at the post-translational level, most likely by decreased assembly of FeS clusters in this cellular compartment. However, the abundance of NBP35, a protein involved in FeS cluster assembly, was similar in *nramp3 nramp4* and wild type. We investigated if aconitase activity could be restored by providing the seedlings with a high concentration of external Fe (200 μM), but the activity was similar to seedlings germinated with 50 μM Fey (Figure 7D). Thus, seedlings are entirely dependent on their vacuolar Fe stores during germination.

**Figure 7.**
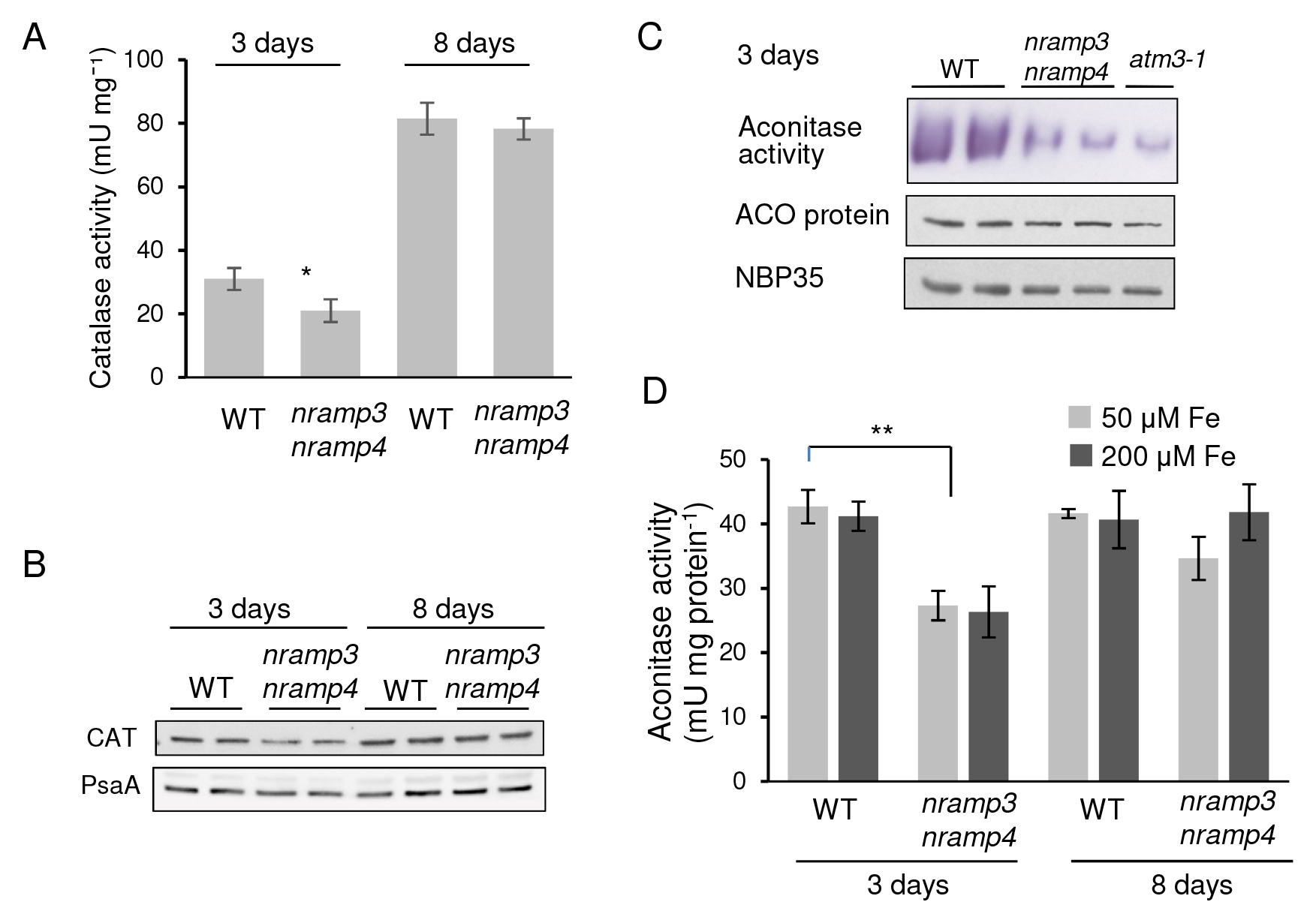
Activities of cytosolic Fe enzymes in *nramp3 nramp4*.
A, Catalase activity, measured by consumption of H_2_O_2_ in a spectrophotometric assay at 240 nm of 3-day and 8-day-old plants. Values represent the mean ± SD (n = 3 – 4), ^*^*p* < 0.05 (unpaired Student's t-test). B, Catalase protein detected by Western bot analysis. The membrane was reprobed with antibodies against PsaA to show equal protein loading. C, Ingel staining of aconitase activity in 3-day-old WT and *nrampS nramp4* seedlings (top panel). The majority of the activity is attributable to a large cytosolic pool of ACO3, depending on ATM3 for maturation of the FeS cluster (Hooks et al., 2014). The same protein extracts were subjected to Western blot analysis with antibodies against aconitase (ACO) and NBP35. D, Aconitase activity in total cell extracts of WT and *nrampS nramp4* with 50 and 200 μM Fe. Values are the mean ± SD (n = 2–4). ^**^*p* < 0.05 (2-tailed Student’s t-test).

### Cell expansion and nutrient transport are actively restricted during the early stage of *nramp3 nramp4* seedling development

A large proportion of downregulated genes (15 out of 78, Figure 3D and Table S4) encode extracellular or cell wall proteins. Among these were numerous extensin-like proteins (EXT10, EXT12, AT3G54580, AT4G08400 and AT4G08410) as well as pectin methyl esterase (PME5) that allow cell wall extension and have a role in root hair formation. This indicates that failure to mobilize seed Fe stores triggers a transcriptionally regulated growth arrest, and consequently downregulation of cell wall extension. In addition, genes that encode plasma membrane proteins were also downregulated. Among them was *RHS15*, which encodes a protein that is required for root hair development. Moreover, several nutrient transporter genes are down regulated in agreement with a restriction of growth. These include the amino acid transporter AAP2, involved in phloem loading and amino acid distribution to the embryo; YSL1, involved in transport of Fe-chelates (Le Jean & Schikora 2005), the sulfate transporter (SULTR1;1) normally upregulated by sulfur deficiency (Barberon et al., 2008); and the phosphate transporter PHO1 involved in phosphate translocation to shoots (Wege et al., 2016). Interestingly, while Fe deficiency responses were still up at day 8, many genes that were downregulated at day 3, including extensins and nutrient transporters, recovered wild-type levels of expression and ultimately growth was not affected in *nramp3 nramp4* in the conditions used for this analysis.

In conclusion, our data indicate that *nramp3 nramp4* seeds are Fe deficient immediately upon hydration and respond by upregulating Fe-deficiency response genes during germination while they prepare for growth arrest in a coordinated manner. Fe-dependent metabolism in mitochondria was maintained, which is essential to release energy from lipid stores and sustain germination and growth. In contrast, chloroplast genes were downregulated indicating that establishment of autotrophy is not the main priority when Fe is lacking. Delay in the establishment of photosynthesis represents a highly efficient way to spare Fe as chloroplasts are the main sink for Fe in photosynthetically active cells. Interestingly, Fe deficiency responses were sustained even after the seedling was able to acquire Fe from the medium to restore growth and photosynthetic function.

## METHODS

### Plant material and growth

*Arabidopsis thaliana* ecotype Columbia (Col-0) plants were used as the wild type. The T-DNA insertion lines SALK_023049 for *nramp3-2* and SALK_085986 for *nramp4-3* (Figure S1A) were crossed and the *nramp3-2 nramp4-3* double mutant was selected in the F2 generation (Molins et al., 2013). The double mutant is named *nramp3 nramp4* for simplicity throughout this study. Wild-type and mutant plants were grown side-by-side in controlled environment conditions (16 h light / 8 h dark, 22 °C, light intensity of 120 – 160 μmol m^−2^ s^−1^) and seeds from 24 plants from each line were harvested and pooled. Seeds were sterilised using chlorine gas, vernalized for 2 days at 4 °C and germinated in a minimum volume of half-strength Murashige and Skoog liquid medium in a Sanyo Versatile Environmental Test chamber under the standard long-day conditions.

### Protein blot analysis

Protein extracts were separated by SDS-PAGE and transferred under semi-dry conditions to nitrocellulose membrane for immunolabelling. Ponceau-S staining of the membranes was used to confirm equal protein loading and successful transfer. Polyclonal antibodies against Arabidopsis NBP35 and aconitase were as previously described (Bych *et al*., 2008; Bernard *et al*., 2009). Polyclonal antibodies against catalase, ferritin, MnSOD, CuZnSOD, FeSOD, PsbA and PsaA were from Agrisera (Umea, Sweden). NFU4 and NFU5 were detected using polyclonal antibodies against NFU4 which recognize both homologous proteins.

### Enzyme assays

In-gel assays for aconitase were as previously described (Bernard et al., 2009). Catalase activity was measured using a spectrophotometric assay for H_2_O_2_ (Beers et al., 1952). Superoxide dismutase activity was measured according to Chu et al. (2005). Blue native PAGE and in-gel activity assays were completed as previously reported (Sabar et al., 2005). Guaiacol peroxidase activity was determined spectrophotometrically (Molins et al., 2013). For all enzyme assays, activity was normalized to protein concentration in the extract, which was determined using BioRad Protein Assay Dye Reagent. Chlorophylls were extracted using 1 ml acetone from 35 mg tissue and the concentrations were quantified using absorption at 662 nm and 645 nm, as previously reported (Lichtenthaler, 1987). Ferric chelate reductase activity was determined as previously described (Yi et al., 1996), except that whole seedlings were submerged in the assay solution.

### RNA extraction

Time points of 1-day old (imbibed), 3-days old (radical emergence) and 8-days old (cotyledon emergence) plants were harvested for RNA extraction in triplicate for wild type (Col 0) and *nramp3 nramp4* (18 samples). RNA from imbibed seeds was isolated as described in Penfield et al., (2005) with minor modifications. In brief, 30–40 mg of flash frozen seed (based on wet seed weight) were ground with a mini-pestle in 300 |il chilled XT buffer (0.2 M sodium borate, 30 mM EGTA, 1% (w/v) SDS, 1% (w/v) sodium deoxycholate, 2% (w/v) polyvinylpyrollidone, 10 mM DTT, and 1% (w/v) IGEPAL [pH 9.0]) treated with diethyl pyrocarbonate. After thawing, 12 μl proteinase K was added and the mixture was incubated at 42 °C for 90 min, followed by addition of 24 μl 2 M KCl and 60 min incubation on ice. The supernatant was collected after centrifugation at 4 °C and the RNA was precipitated at −20 °C for 2 hr (or overnight) with 108 μl 8 M LiCl. The RNA was collected by centrifugation at 4 °C and redissolved in 30 μl RNase-free water. The RNA was purified using a DNase I kit (Promega) and the RNeasy Plant Mini kit (Qiagen), starting with the addition of 60 μl RNase-free water and 350 μl RLT buffer. Extraction of RNA from seeds with radical emergence or cotyledon growth was completed using the RNeasy Plant Mini kit (Qiagen). Concentration of total RNA was measured using a NanoDrop 1000 Spectrophotometer (Thermo Scientific).

### RNA-sequencing

Adequate quality of the RNA for RNA-sequencing was verified using a Bioanalyser 2100 (Agilent). Library preparation and RNA-sequencing were performed by Oxford Gene Technology (Begbroke, UK). RNA libraries were prepared using an Illumina TruSeq Stranded mRNA kit and sequenced using an Illumina HiSeq 2500 with 100 bp paired-end reads. All 18 samples were run in the same lane. The total library size before mapping ranged from 29–47 million reads (Table S1), with an average read count per sample of 8.88 million paired-end reads (100 bp). Read trimming was used to remove adapter sequences.

RNA-sequencing reads were aligned to the *Arabidopsis thaliana* reference genome (TAIR10) using CLC Genomics Workbench using default parameters, except we used a length fraction of 0.7 and similarity fraction of 0.95.

### Normalisation and statistical analysis

Read count data sets were filtered by removing genes with low read counts (counts per million < 2 in at least 4 samples). Normalisation and differential expression was conducted with the edgeR Bioconductor package (McCarthy et al., 2012; Robinson et al., 2010). The library sizes were normalised using the trimmed mean of M-values (TMM) and then statistically analysed using a Negative Binomial Generalised Linear model (GLM), see Table S5. The Benjamini and Hochberg’s algorithm was used to control the false discovery rate (FDR) (Benjamini et al., 1995). To construct the heatmaps Heatmap.2 gplots package (gplots) was used.

### Quantitative reverse transcription polymerase chain reaction (qRT-PCR)

For each sample, 2.4 μg of total RNA was depleted of genomic DNA contamination using TurboDNAse (Ambion), and reverse transcribed to cDNA using Superscript III (Thermo). RT-qPCR reactions were made using SensiFAST master-mix (Bioline), in 20 μl volumes, each with 20 ng of cDNA. Reactions were measured in a Bio-Rad CFX-96 real-time PCR system and cycled as per the Bioline protocol. Data were analysed using the Bio-Rad CFX Manager 3.1 software, and were normalised using primer efficiency. All data points are from 3 independent biological replicates, measured in three technical replicates (n = 9). The house keeping genes *SAND* (*AT2G28390*) and *TIP41-like (AT4G34270*) were used as reference genes, as they are unaffected by Fe levels in *A. thaliana* (Han *et al.*, 2013). See Table S6 for primer sequences.

## ACKNOWLEDGEMENTS

The authors would like to acknowledge the following sources of funding: Biotechnology and Biological Sciences Research Council (BB/K008838/1 to E.L.B. and JB; BBSRC institute strategic programmes grants (BB/P012523/1, BB/P012574/1) to R.T.G. and J.B. The John Innes Foundation to A.E.M., Agence Nationale de la Recherche grants ANR-16-CE20-0019-2 to S.T.; ANR-13-ADAP-0004 and LabEx Saclay Plant Sciences-SPS (ANR-10-LABX-0040-SPS) to S.M. Antibodies against NFU4 were from Nicolas Rouhier (University of Lorraine). We also wish to thank Anja Krieger Liszkay for helpful discussion and sharing antibodies against PsaA and PsbA, and Jorge Rodriguez-Celma for valuable comments to the manuscript.

## Author contributions

E.L.B, V.S.G.T., A.E.M., R.T.G. performed the experiments; E.L.B., V.S.G.T., S.M. analysed the RNA-seq data set; S.T. and J.B. conceived the project; E.L.B, S.T. and J.B. wrote the article with contributions of the other authors.

---

**Figure S1.**
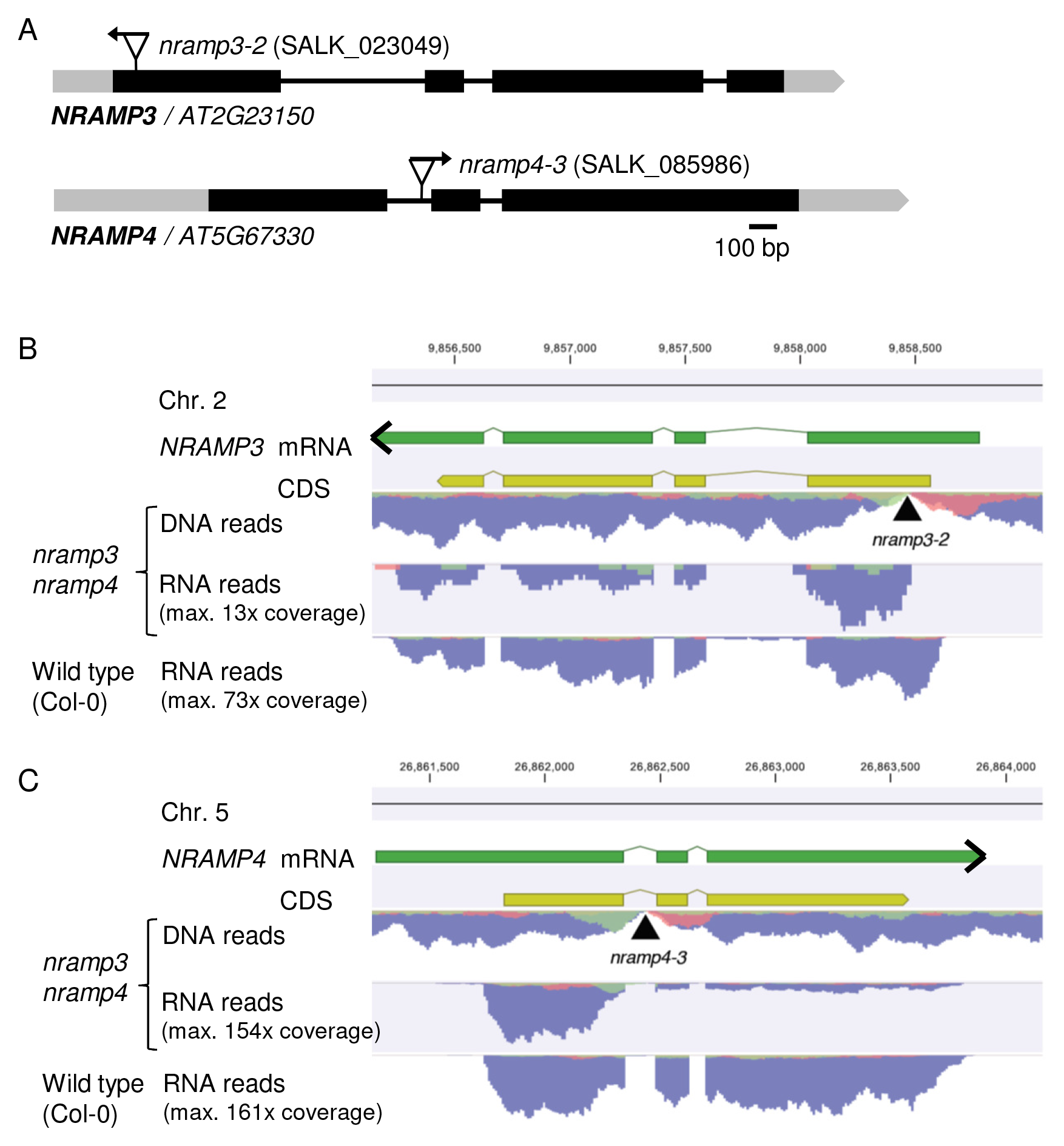
Sequence analysis of the *nramp3-2 nramp4-3* double mutant. A, Gene models of Arabidopsis *NRAMP3 (AT2G23150*) and *NRAMP4 (AT5G67330*) showing the sites of T-DNA insertions for the *nramp3-2 and nramp4-3* mutant alleles. Arrows indicate the left border of the T-DNA. B and C, DNA and RNA sequencing reads from the *nramp3-2 nramp4-3* double mutant and wild type mapped to *NRAMP3* (B) and *NRAMP4* (C). Reads were mapped to the TAIR10 Arabidopsis genome sequence. Blue represents paired reads, red and green represent forward and reverse non-paired reads. The mapped DNA reads reveal the position of the T-DNA insertions (black triangles). Analysis of the mapped RNA reads shows that in the *nramp3 nramp4* mutant transcription of *NRAMP3* starts after the T-DNA insertion and transcripts thus lack the ATG start codon. The number of full-length *NRAMP4* transcripts is significantly reduced in the *nramp4-3* allele.

**Figure S2.**
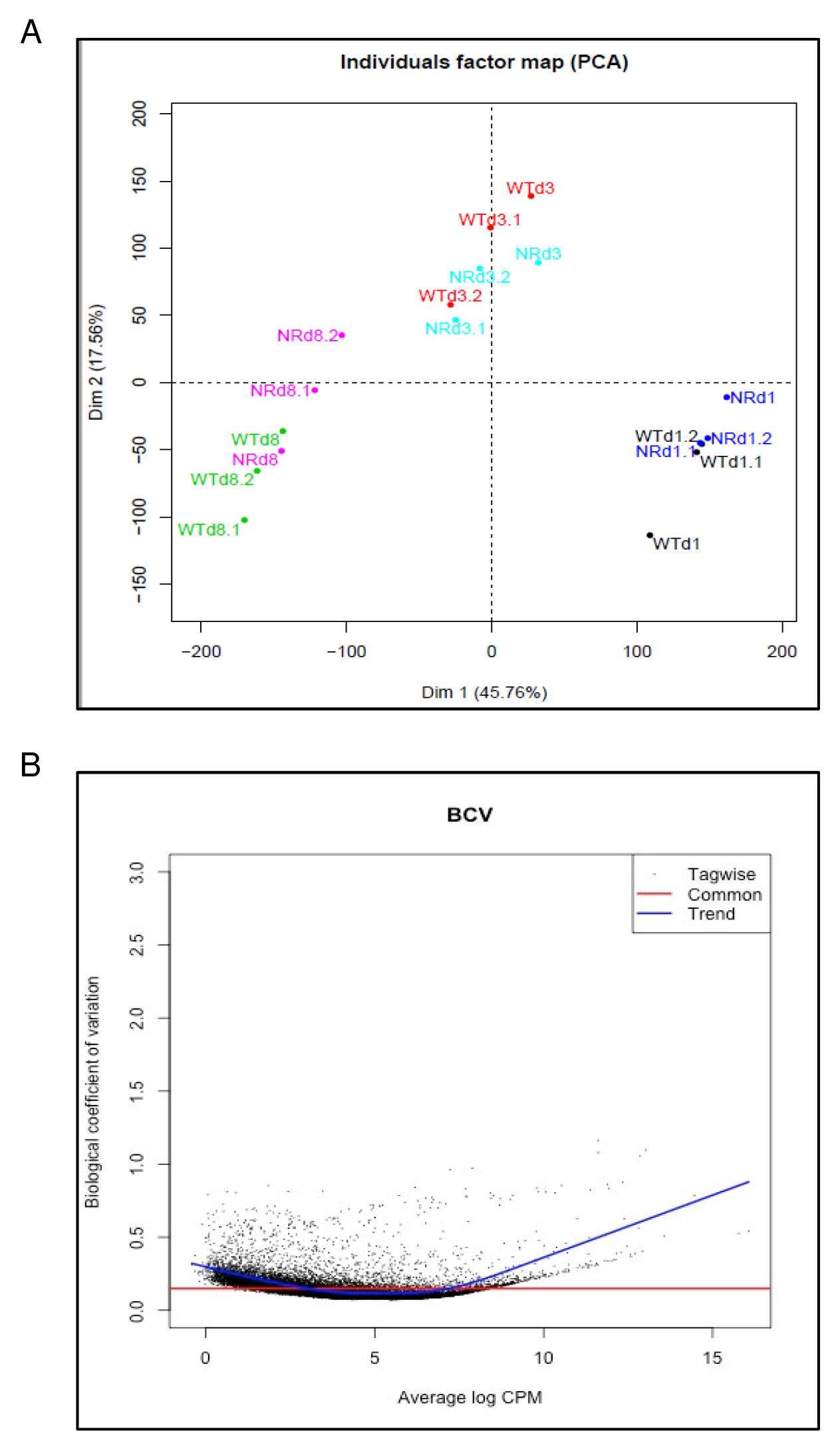
Quality of the sequencing data. A, Multi-dimensional scaling plot (MDS) of the principle component analysis (PCA) to examine the samples for outliers. B, Biological coefficient of variation (BCV) to identify variation of gene expression between replicate RNA samples. Dispersion: 0.02221, BCV: 0.149.

**Table S1.** Percentage of paired reads that were mapped to transcripts

|  | **Total number of reads**<br>**(average of replicates)** | **Reads mapped in pairs**<br>**or broken pairs (%)** |
| --- | --- | --- |
| Wild type_day 1 | 37,045,051 | 85 |
| Wild type_day 3 | 41,064,517 | 91 |
| Wild type_day 8 | 35,679,516 | 84 |
| *nramp3 nramp4*_day 1 | 36,266,804 | 91 |
| *nramp3 nramp4*_day 3 | 35,732,362 | 90 |
| *Nramp3 nramp4*_day 8 | 39,470,477 | 89 |

**Table S2.** Number of differentially expressed genes with >3-fold change (*P* < 0.05). Total number of genes analysed was 18,493.

|  | Day 1 | Day 3 | Day 8 |
| --- | --- | --- | --- |
| Up | 13 | 78 | 135 |
| 0 | 18473 | 18376 | 18295 |
| Down | 7 | 39 | 63 |

**Table S3.** Genes UPregulated in 3-day-old *nramp3nramp4* compared to wild type. Expression of genes upregulated in 3-day-old plants were compared between the 1-day-old and 8-day-old timepoints.

|  |  |  | day 1 | day 3 | day 8 |
| --- | --- | --- | --- | --- | --- |
| ID | Symbol | Description | FC (log2) | FC (log2) | FC (log2) |
| AT1G07367 | ncRNA | ncRNA | 4.6 | 5.8 | 2.3 |
| AT1G13609 | IRP4 | Defensin-like (DEFL) family protein | 3.4 | 5.5 | 2.3 |
| AT1G47395 | IRP2 | hypothetical protein | 2.7 | 5.7 | 1.9 |
| AT1G47400 | IRP1 | hypothetical protein | 3.0 | 5.7 | 2.5 |
| AT1G62420 |  | DUF506 family protein (DUF506) | 0.3 | 3.2 | -0.5 |
| AT1G74770 | BTSL1 | zinc ion binding protein | 2.1 | 3.1 | 0.8 |
| AT1G75945 |  | hypothetical protein | 6.2 | 7.6 | 7.3 |
| AT2G01422 |  | unknown | 2.7 | 3.9 | 3.3 |
| AT2G30760 |  | hypothetical protein | 1.9 | 3.5 | 1.8 |
| AT2G30766 |  | hypothetical protein | 1.7 | 3.6 | 0.5 |
| AT2G41730 |  | calcium-binding site protein | 0.3 | 3.4 | -0.2 |
| AT2G42750 | DJC77 | DNAJ heat shock N-terminal domain-containing protein | 0.5 | 2.2 | 0.2 |
| AT3G18290 | BTS | zinc finger protein-like protein | 0.9 | 2.5 | 0.5 |
| AT4G05275 |  | ncRNA | 13.1 | 12.7 | 12.0 |
| AT4G08867 |  | hypothetical protein | 2.4 | 6.0 | 4.4 |
| AT4G08869 |  | defensin-like (DEFL) family protein | 1.8 | 6.3 | 3.3 |
| AT4G16370 | OPT3 | oligopeptide transporter | 1.9 | 2.6 | 0.5 |
| AT4G36700 |  | RmlC-like cupins superfamily protein | 0.1 | 3.6 | 2.4 |
| AT4G37370 | CYP81D8 | cytochrome p450, family 81, subfamily d, polypeptide 8 | -0.2 | 2.7 | -1.1 |
| AT5G04150 | BHLH101 | basic helix-loop-helix (bHLH) DNA-binding superfamily protein | 2.5 | 5.4 | 2.5 |
| AT5G05250 | IRP6 | hypothetical protein | 1.1 | 2.8 | 2.2 |
| AT5G25840 |  | DUF1677 family protein | 1.5 | 2.8 | -0.9 |
| AT5G33390 |  | glycine-rich protein | 2.7 | 3.0 | 3.8 |
| AT5G67370 | CGLD27 | DUF1230 family protein | 2.1 | 3.7 | 0.6 |
| AT5G20230 | SAG14 | blue-copper-binding protein | 0.2 | 3.8 | 0.3 |
| AT2G41240 | BHLH100 | basic helix-loop-helix protein 100 | 4.6 | 7.2 | 1.8 |
| AT1G64980 | CDI | Nucleotide-diphospho-sugar transferases superfamily protein | 1.1 | 2.1 | 0.5 |
| AT1G03990 | FAO1 | Long-chain fatty alcohol dehydrogenase family protein | -0.1 | 2.4 | -0.4 |
| AT1G01580 | FRO2 | ferric reduction oxidase 2 | 0.2 | 2.4 | 4.2 |
| AT1G23020 | FRO3 | ferric reduction oxidase 3 | 1.6 | 2.2 | 0.9 |
| AT2G02010 | GAD4 | glutamate decarboxylase 4 | -3.8 | 4.4 | -2.1 |
| AT4G19690 | IRT1 | iron-regulated transporter 1 | -3.5 | 6.8 | 5.1 |
| AT1G03470 | NET3A | networked 3a | 0.0 | 2.0 | 0.8 |
| AT3G56970 | BHLH038 | basic helix-loop-helix (bHLH) DNA-binding superfamily protein | 3.8 | 6.7 | 2.1 |
| AT3G56980 | BHLH039 | basic helix-loop-helix (bHLH) DNA-binding superfamily protein | 3.3 | 6.0 | 1.7 |
| AT5G53450 | ORG1 | OBP3-responsive protein 1 | 1.4 | 3.4 | 1.1 |
| AT1G14550 |  | Peroxidase superfamily protein | -2.6 | 3.5 | -7.8 |
| AT1G05675 | UGT74E1 | UDP-glycosyltransferase 74E1 | 0.6 | 4.5 | -1.3 |
| AT5G13740 | ZIF1 | zinc induced facilitator 1 | 1.2 | 3.1 | 1.0 |

**Table S4.**
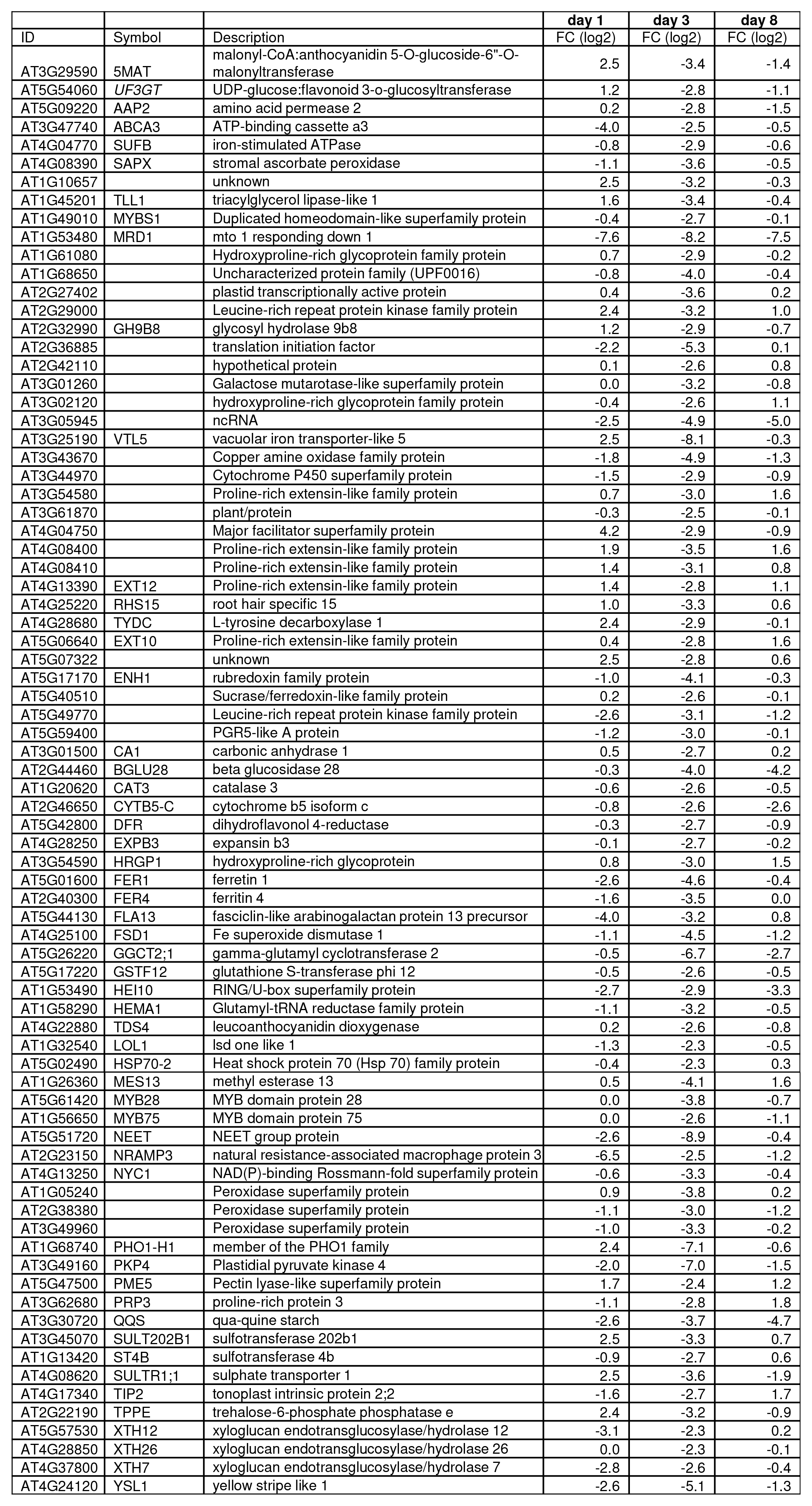
Genes DOWNregulated in 3-day-old *nramp3nramp4* compared to wild type. Expression of genes upregulated in 3-day-old plants were compared between the 1-day-old and 8-day-old timepoints.

**Table S5.** Normalisation factors calculated using TMM

|  | Group library size | Normalisation factor |
| --- | --- | --- |
| WTd1.1 | 29968889 | 0.97655508 |
| WTd1.2 | 28933408 | 1.29494176 |
| WTd1.3 | 33548999 | 1.14622239 |
| NRd1.1 | 36247600 | 1.1973107 |
| NRd1.2 | 31038949 | 1.22628428 |
| NRd1.3 | 32174294 | 1.2102143 |
| WTd3.1 | 41908877 | 1.1327056 |
| WTd3.2 | 37553980 | 1.11334062 |
| WTd3.3 | 32331672 | 1.02069473 |
| NRd3.1 | 34528796 | 1.14111771 |
| NRd3.2 | 28459074 | 1.10606036 |
| NRd3.3 | 33039728 | 1.11082451 |
| WTd8.1 | 32061776 | 0.81591707 |
| WTd8.2 | 25166347 | 0.79472869 |
| WTd8.3 | 32300798 | 0.72914147 |
| NRd8.1 | 30350879 | 0.75091096 |
| NRd8.2 | 34070193 | 0.79860716 |
| NRd8.3 | 41037330 | 0.75880429 |

**Table S6.**
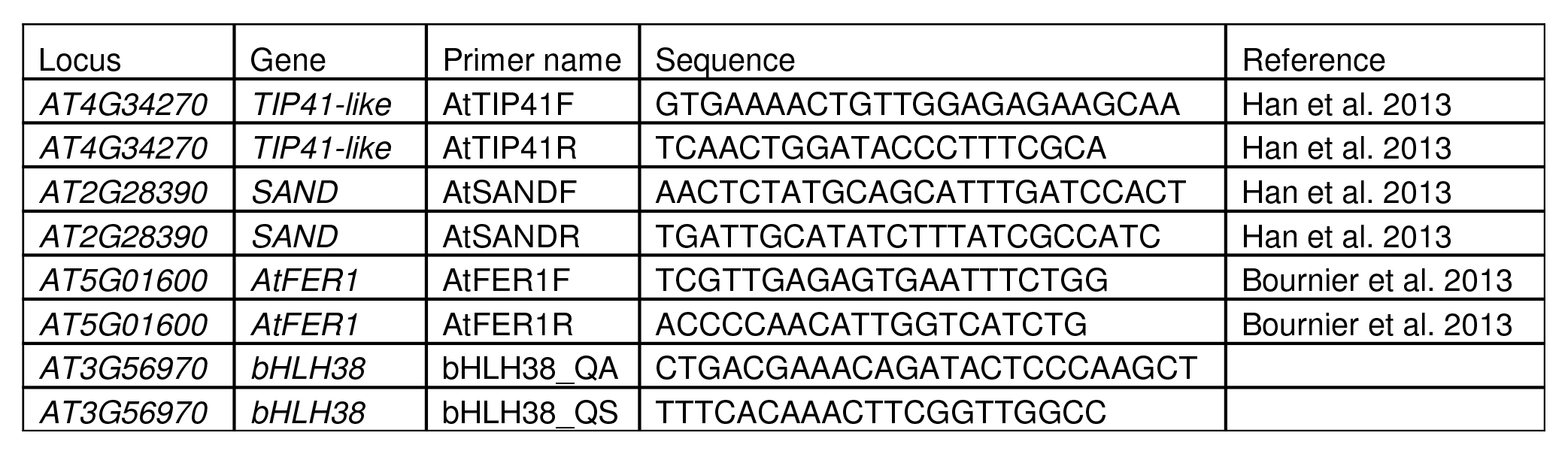
Primers used in qRT-PCR

